# A Novel Tool for Multi-Omics Network Integration and Visualization: A Study of Glioma Heterogeneity

**DOI:** 10.1101/2024.03.18.585500

**Authors:** Roberta Coletti, João Carrilho, Eduarda P. Martins, Céline S. Gonçalves, Bruno M. Costa, Marta B. Lopes

**Affiliations:** Center for Mathematics and Applications (NOVA Math), NOVA School of Science and Technology, Largo da Torre, Caparica, 2825-149, Portugal; NOVA School of Science and Technology, NOVA University of Lisbon, Largo da Torre, Caparica, 2825-149, Portugal; Life and Health Sciences Research Institute (ICVS), School of Medicine, University of Minho, Campus Gualtar, Braga, 4710-057, Portugal; ICVS/3B’s - PT Government Associate Laboratory, Braga/Guimarães, Portugal; UNIDEMI, Department of Mechanical and Industrial Engineering, NOVA School of Science and Technology, Largo da Torre, Caparica, 2825-149, Portugal

**Keywords:** MINGLE, network, multi-omics, glioma, GBM, graph, visualization, variable selection, biomarkers

## Abstract

Gliomas are highly heterogeneous tumors with generally poor prognoses. Leveraging multi-omics data and network analysis holds great promise in uncovering crucial signatures and molecular relationships that elucidate glioma heterogeneity. However, the complexity of the problem and the high dimensionality of the data increase the challenges of integrating information across various biological levels. In this study, we developed a framework comprising two steps for variable selection based on sparse network estimation from various omics. Subsequently, we introduced MINGLE (Multi-omics Integrated Network for GraphicaL Exploration), a novel methodology designed to merge distinct multi-omics information into a single network, enabling the identification of underlying relations through an innovative integrated visualization. Applying this method to glioma data, with patients grouped according to the newest glioma classification guidelines, led to the selection of variables as potential candidates for novel glioma-type-specific biomarkers.

## 1. Introduction

Gliomas are tumors originating from glial or stem cells in the brain, which exhibit a remarkable level of heterogeneity, both at the molecular and histo-pathological levels [1]. This inherent complexity has presented significant challenges in glioma classification and understanding, ultimately impacting treatment decisions and patient outcomes [2]. The advent of modern techniques for data acquisition allowed the generation of huge amounts of data describing different molecular entities within a biological system, i.e., omics data [3]. The analysis of this type of data substantially increased the opportunities to disclose important molecular insights for the different glioma types, contributing to improve patient stratification [4, 5]. As a result, the World Health Organization (WHO) recently updated the standards for the classification of Tumors of the Central Nervous System (CNS), releasing the last edition in 2021 (2021-WHO) [6], ultimately distinguishing three main adult-type diffuse gliomas: astrocytoma, oligodendroglioma, and glioblastoma (GBM). These advances have also contributed to the development of modern therapies (such as immunotherapy), which currently complement the classical approaches (chemotherapy, surgery, and radiation) [7]. Despite this, glioma patients often show poor overall survival [8], indicating that further efforts are needed to improve the life expectancy and quality of glioma patients.

In this context, mathematical and statistical methods might be extremely useful. In particular, graphical models might deal with the estimation of networks of molecular features at different biological levels, allowing the disclosure of potentially important relations between multiple molecular entities and clinical outcomes. Nevertheless, traditional approaches fail when dealing with high-dimensional data, when the number of variables (p) vastly outnumbers the number of samples (n), as it occurs with omics data [9]. Dimensionality reduction techniques can address this issue, reducing the number of features while maintaining the overall information. In the past years, several methods have been proposed to infer sparse networks from biological data, with a particular focus on gene regulatory networks based on transcriptomics data [10]. However, the incorporation of multiple molecular levels through multi-omics integration leads to a more comprehensive perspective on the complex molecular landscape regulating cell behaviour [11]. The multi-omics approach enables researchers and clinicians to identify subtle patterns and associations unfeasible to discern by considering a single omics view [12, 13, 14]. On the other hand, this integration might be challenging due to the high dimensionality and heterogeneity of these data [15].

Here, we propose a novel methodology specifically designed to define a Multi-omics Integrated Network for GraphicaL Exploration (MINGLE). Despite the method being applicable to different omics, here we consider RNA– Sequencing data (*transcriptomics*), and DNA–methylation data (*methylomics*) of patients from The Cancer Genome Atlas (TCGA) glioma projects (namely, TCGA-LGG, and TCGA-GBM), previously updated according to the 2021-WHO diagnostic annotations [16]. The selection of these two omics was motivated by the close relationship existing between them. Indeed, whereas RNA sequencing drives all cell processes, methylation of small regions of DNA might affect transcript reading [17], significantly impacting cancer regulation [18]. In particular, in the context of glioma, methylation might explain the pathogenesis and clinical behavior of the different cancer types [19, 20], therefore being promising information to include in a multi-omics study aiming at understanding glioma heterogeneity. The designed pipeline is constituted by two sequential steps of network-based variable selection achieved by graphical models. Specifically, we applied Graphical Lasso [21] and Joint Graphical Lasso [22] to estimate sparse networks from each omics layer, by focusing the analysis on the connected nodes. The estimated undirected relations are then used to merge the information obtained from the two omics into a single integrated graph by MINGLE. This new representation leads to a network of genes whose relations are based on methylomics data, allowing the discovery of underlying gene relations impossible to detect from a single-omics study.

The analysis here reported is focused on GBM, as it represents the most aggressive and most frequent malignant glioma, with a very poor survival. Disclosing underlying molecular relations potentially associated with GBM evolution is extremely important to identify novel therapeutic targets, design innovative and more effective therapies, and ultimately improve patient survival and quality of life. Nevertheless, equivalent analyses can be easily performed on different glioma types through the provided R codes and tables, which allow the independent exploration of our results. These materials will be made available after publication. To validate the provided outcomes, a mathematical validation of the networks inferred was performed, followed by classification to formally evaluate the ability of the selected features to distinguish among the glioma types. Importantly, survival analysis on a subset of variables was also applied to assess the prognostic value of these features.

## 2. Materials and methods

### 2.1. Network inference and variable selection

Let 𝔾 be a graph with *p* nodes ***X*** = (*X*_1_, …, *X*_*p*_) having a multivariate normal distribution, and with theoretical and empirical covariance matrix Σ and *S*, respectively. The edges of the graph 𝔾 can be estimated based on the non-zero entrances of Θ = Σ^*-*1^, also known as precision matrix [23]. In particular, each entrance *θ*_*i,j*_ describes the conditional dependence between the variables *X*_*i*_ and *X*_*j*_, given the others *X*_*k*_, *k* = 1, …, *p, k* ≠ *i, j*.

Given a dataset with *n* observation of *X*_1_, …, *X*_*p*_ variables, if *p* » *n* estimating the fully-connected precision matrix is unfeasible, and methods leading to sparse networks should be employed. The graphical lasso (*glasso*) [21] finds sparse Σ and Θ matrices, by solving the following Gaussian log-likelihood maximization problem:

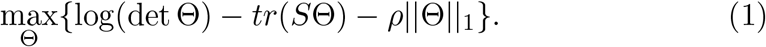

The presence of the regularization term *ρ*||Θ||_1_ makes the solution sparse, which degree is regulated by the parameter *ρ*.

Let 𝔾_1_, …, 𝔾_*k*_ be *k* graphs with the same nodes ***X*** = (*X*_1_, …, *X*_*p*_), and different sets of edges, represented by the precision matrices **Θ** = (Θ^(1)^, …, Θ^(*k*)^). The Joint Graphical Lasso (JGL) methods [22] allows the joint estimation of *k* precision matrices, by a generalization the function (1):

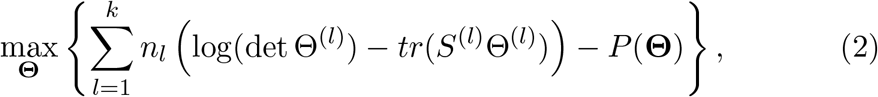

where *n*_*l*_ and *S*^(*l*)^, for *l* = 1, …, *k* are the number of observations of the *l-*th dataset, and the corresponding empirical covariance matrices. The penalization term *P* (**Θ**) affects the sparsity of the estimated **Θ**; in this work, we employed the Fused Graphical Lasso penalty:

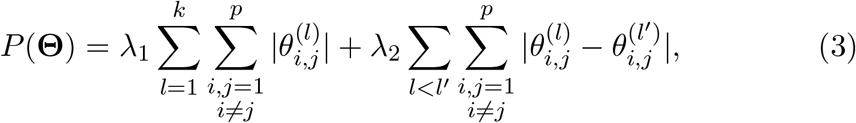

where the parameter *λ*_1_ regulates the amount of sparsity (as *ρ* in glasso equation (1)), while *λ*_2_ encourages similarity across edges.

The sparsity of the precision matrix determines a network-based variable selection. Indeed, *θ*_*i,j*_ = 0, for a certain *i* and *j* (*i* ≠ *j*), means that the variables *X*_*i*_ and *X*_*j*_ are conditionally independent given the others *X*_*k*_, *k* = 1, …, *p, k* ≠ *i, j*. In highly-sparse networks, a certain number of variables will appear as isolated nodes, implying that the corresponding features could be removed, as they do not impact on the overall network.

### 2.2. Sparse multinomial logistic regression with the elastic net penalty

Let ***X*** = (*X*_1_, …, *X*_*p*_) be a set of *p* variables among which *n* observations are collected. Let we assume that each observation is attributed with a categorical variable ***C***, with *L* possible levels. Zhu and Hastie [24] proposed a symmetrical approach for the estimation of the probabilities of a given sample being attributed to level *ℓ* = 1,. .., *L*:

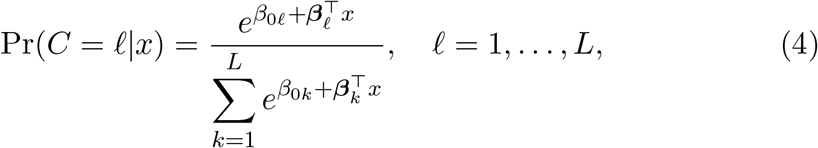

where ***β***_*ℓ*_ = (*β*_1*ℓ*_,. .., *β*_*pℓ*_). The coefficients *β*_0*ℓ*_ and ***β***_*ℓ*_ are then estimated by solving the following minimization problem:

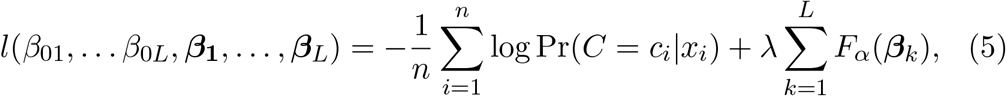

where *n* represents the number of observations, *x*_*i*_ the *i*-th observation, *c*_*i*_ the class which *x*_*i*_ is attributed to and *λ* the strength of the penalty. *F*_*α*_ represents the elastic net penalty function, defined as

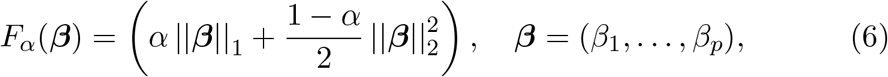

with *α* controlling the balance between the Lasso and Ridge penalties and *λ* controlling the strength of the penalty. The regularization term induces sparsity in the estimated coefficients *α* _≠_, allowing variable selection.

### 2.3. Dataset preprocessing

The glioma datasets are publicly available from the TCGA portal, under the projects TCGA-LGG and TCGA-GBM [25, 26, 27]. To update the glioma types according to the latest 2021 WHO classification of brain tumors [6], we followed the methodology described in our previous work [16], integrating the samples’ molecular profiles [28]. We considered only patients included in both omics datasets (transcriptomics and methylomics), and having survival time information, to perform survival analysis on a subset of selected features. After this preprocessing, the samples were grouped in three datasets constituted by 249 astrocytoma, 166 oligodendroglioma and 112 GBM samples.

Before the application of the network inference methods, the variables of each glioma-type dataset were normalized through the *huge.npn* function from *huge* R package [29], which applies a non-parametric normal (nonparanormal) transformation aiming to obtain variables with multivariate Gaussian distribution. For our analysis, we considered only variables normally distributed according to the Jarque-Bera test [30].

The TCGA transcriptomics datasets are constituted by a matrix containing the expression of 20501 genes, already normalized by Transcripts Per Million (TPM) and upper quartile normalizations. After the nonparanormal normalization, a dataset of 16654 normally distributed genes was obtained. The complete TCGA methylomics datasets contain 485577 variables, expressing the ratio of methylation of each gene’s methylation site. To reduce the dimensionality, after removing variables with zero variance and non-valid entries, filters based on previous knowledge were applied. In particular, we only considered C-phosphate-G (CpG) sites, as more than 98% of DNA methylation occurs in a CpG dinucleotide [31], and we excluded from the analysis: probes related to sexual chromosomes [32], cross-reactive and polymorphic CpGs, as suggested in [33]. At the end of this procedure, methylation datasets contain 340427 variables. Given that the network inference methods are unable to deal with this high dimensionality, we developed a framework to integrate the two omics, while reducing the dimensionality of this dataset, as explained in the next section.

### 2.4. Pipeline and implementation

The general framework and the integrated multi-omics strategy developed for this study is schematized in Figure 1. As a first step after data preprocessing, JGL was applied to the RNA-sequencing dataset, by considering three classes corresponding to the main glioma types (astrocytoma, oligodendroglioma, and GBM), through the `JGL` function from the same package [34]. We tested several combinations of parameters (Table S1), yet only the results obtained by setting *λ*_1_ = 0.90 and *λ*_2_ = 0.001 were explored. JGL determined a selection of 853 variables, specifically 517, 611, and 300 for astrocytoma, oligodendroglioma, and GBM, respectively. To explore the differences among the three glioma types, we focused the analysis on the genes involved in exclusive edges (i.e., edges estimated for a single glioma type), which we assumed to be more representative of the corresponding disease condition. Therefore, in the second step, the subsets of these exclusive pairs of genes were considered, and associated with the corresponding CpG sites in the DNA-methylation dataset by the function `data` from *methylGSA* R package [35]. This association led to three datasets comprising 6650, 6811 and 2917 variables for astrocytoma, oligodendroglioma, and GBM, respectively. Given the high number of variables, we performed another variable selection through *glasso* algorithm, by the `huge.glasso` function from *huge* R package [29] with *ρ* = 0.85, which led to an interpretable amount of features, yet sufficiently large for a comprehensive analysis (Table S2).

**Figure 1:**
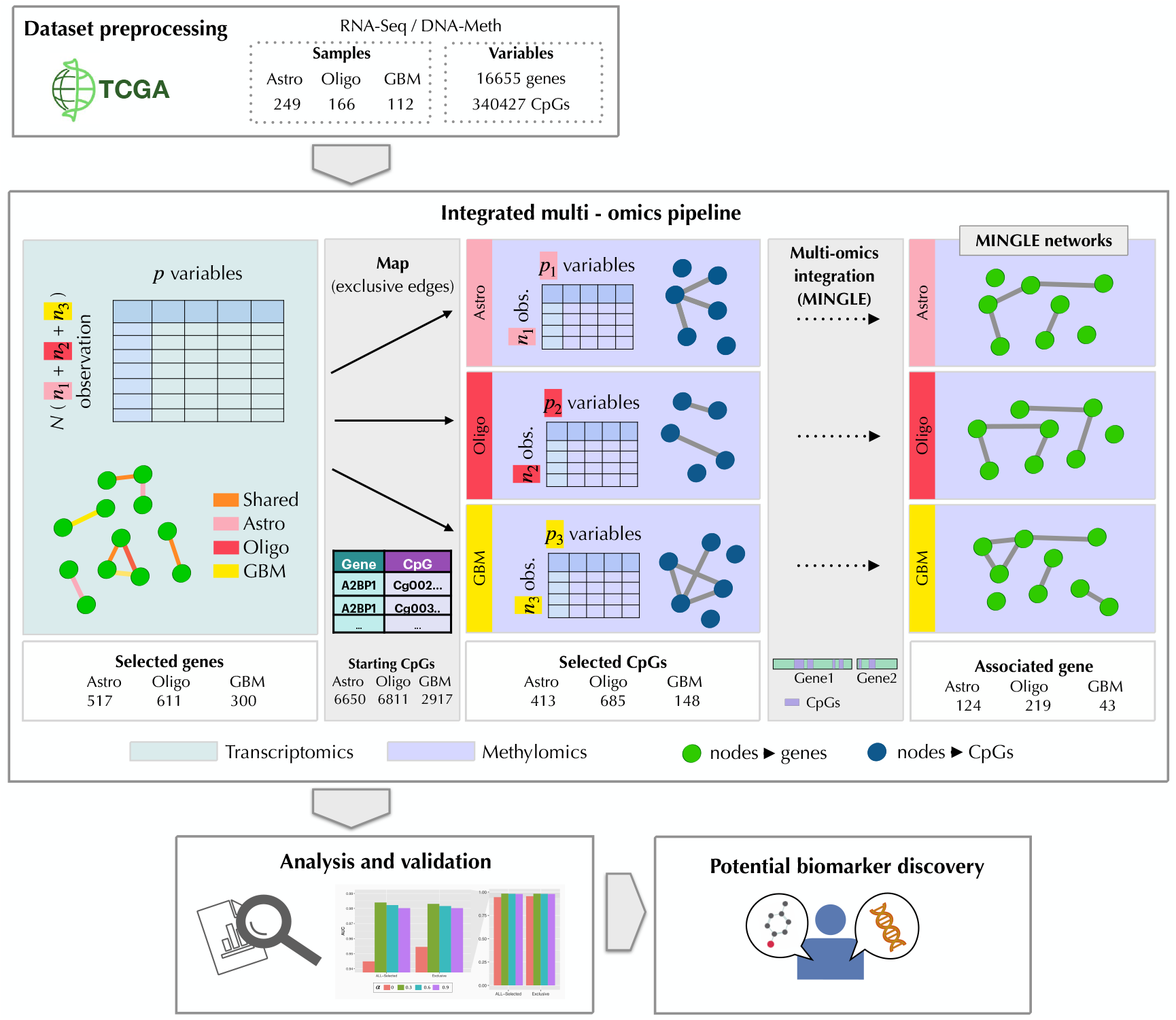
Workflow of the analysis and multi-omics integration scheme. After data loading and preprocessing, the obtained datasets were used as input for JGL and glasso algorithm. In the integrated multi-omics pipeline, green and purple panels refer to networks obtained from transcriptomics and methylomics data, respectively. The nodes are colored differently to stress the molecular entities they represents, i.e., green nodes are genes, while blue nodes are CpG sites. We started our analysis from transcriptomics layer, estimating joint gene networks for the three glioma types. Therefore, only the genes linked by exclusive edges were further analyzed, mapping each one with the respective CpG sites. Methylomics data were divided for the three glioma types, and the network estimation was performed separately (thus, three distinct graphs are obtained). These final outcomes were used to define the MINGLE networks, i.e., networks of genes whose edges are computed based on the estimated methylomics relations. The obtained results were then analyzed and validated to lead to a list of potential biomarkers. Abbreviations: RNA-Seq, RNA sequencing; DNA-Meth, DNA methylation; Astro, astrocytoma; Oligo, oligodendroglioma; GBM, glioblastoma; CpG, CpG site.

The results provided by the two omics were integrated through MINGLE, which defines new gene-networks based on the estimated methylomics networks (see Section 2.6 for details).

The reliability of JGL and glasso results were tested by performing a mathematical validation, as explained in Supplementary Matherial S2. Also, sparse multinomial logistic regression and survival analysis models were employed to investigate the relevance of the biological information carried by the sets of the network-based selected variables. Classification was executed by the `glmnet` R function [36], performing sparse multinomial logistic regression with the elastic net penalty. Different values of the parameter *α ∈* [0, 1] were tested, while the optimal value for *λ* was estimated through 10-fold cross-validation. For each parameter choice, 100 runs were computed, where the dataset was split into 75%-train and 25%-test sets. Classification performances were evaluated by computing the Area under the ROC Curve (AUC). The complete results are resumed in Tables S5 and S6. A subset of the variables detected by our integrated multi-omics study was further investigated by survival analysis, to determine their impact on the patient’s prognosis. To this aim, the univariate Kaplan-Meier method was used, allowing for the estimation of the time of death in a setting of incomplete observations [37]. In particular, for each variable, patients were first stratified into high- and low-risk groups by applying two functions from the `survminer` R package [38]: `surv cutpoint`, which determines the best cut point to divide the two curves, and `surv categorize`, which separates the samples in the two groups. Then, the `survfit` function from the `survminer` R package was employed to estimate the survival probability. This output was used to plot the Kaplan-Meier curves of the statistically significant variables according to the log-rank p-value by the `ggsurvplot` function from the same package.

The codes for reproducing the analysis with personalized choices of regularization parameters will be made available after publication.

### 2.5. Analysis of the results

The first two steps of the proposed workflow lead to a subset of features (genes or CpGs) that are selected based on the estimated networks. The main goal of our analysis is to identify, among these features, the ones having a key role in the disease network, which could be potential biomarkers, specific for a glioma type. We focused our analysis on GBM, but the same study for the other glioma types can be independently explored and reproduced by the provided files and codes.

Given the biological differences between transcriptomics and methylomics layers, we developed different strategies for each step of our pipeline.

Concerning the transcriptomics results, we first evaluated the joint networks to detect subnetworks of interest for deeper analysis in light of shared or exclusive relations among the glioma types. We focused on each subnetwork to disclose relations that could be potentially relevant for GBM.

From the methylomics layer, the glasso algorithm returns a sparse precision matrix for each glioma type. Being that the starting DNA methylation datasets were built to further explore the genes involved in the exclusive relations, each resulting methylomics network is related to a single glioma type. The analysis of these networks cannot be performed as it was done for transcriptomics, given the complexity of the interpretation of such relations. Indeed, genes usually have many regions in which methylation might occur, which could result in a network in which several links are related to the same gene, and affecting the overall understanding and interpretability. To avoid this bias towards the identification of gene-specific methylation subnetworks, and to integrate the results from the two omics, we defined new networks of genes based on methylomics outcomes by MINGLE.

### 2.6. Multi-omics Integrated Network for GraphicaL Exploration (MINGLE)

MINGLE networks are designed to integrate the multi-omics information obtained from the defined two-steps pipeline. In particular, the nodes are defined by the genes associated with the selected CpG sites, while the edges are computed based on the precision matrix estimated by glasso on DNA-Methylation data, allowing the exploration of gene relations based on methylomics relations.

We defined two measures for computing the network edges, leading to different MINGLE networks: one based on the strength of the connections (Strength MINGLE network, *S*_*M*_), and another based on the number of connections (Degree MINGLE network, *D*_*M*_), schematized in Figure 2. Let *g*_*i*_ and *g*_*j*_ be two genes associated with certain sets of selected CpG sites *M*_*i*_ and *M*_*j*_, respectively. To develop the *S*_*M*_, we computed the edge connecting the genes *i* and *j* as:

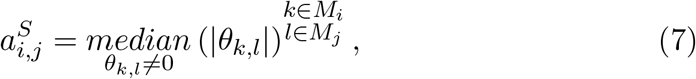

where *k* and *l* are the indices corresponding to all the CpG sites associated with the genes *i* and *j*, respectively.

To define the *D*_*M*_, we considered a weighted degree measure, computing the edge connecting two genes *g*_*i*_ and *g*_*j*_ as:

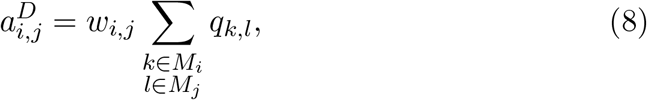

where *q*_*k,l*_ = 1 if *θ*_*k,l*_ ≠ 0, and *q*_*k,l*_ = 0 otherwise. The weights 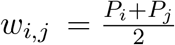, are calculated based on the percentages of selected CpG sites for each gene (*P*_*i*_ and *P*_*j*_). The presence of the weights avoids the edges from being dependent on the different availability of CpG sites per gene, as genes with many CpG sites would naturally have more probability to appear highly connected, introducing a bias in the computed edges.

**Figure 2:**
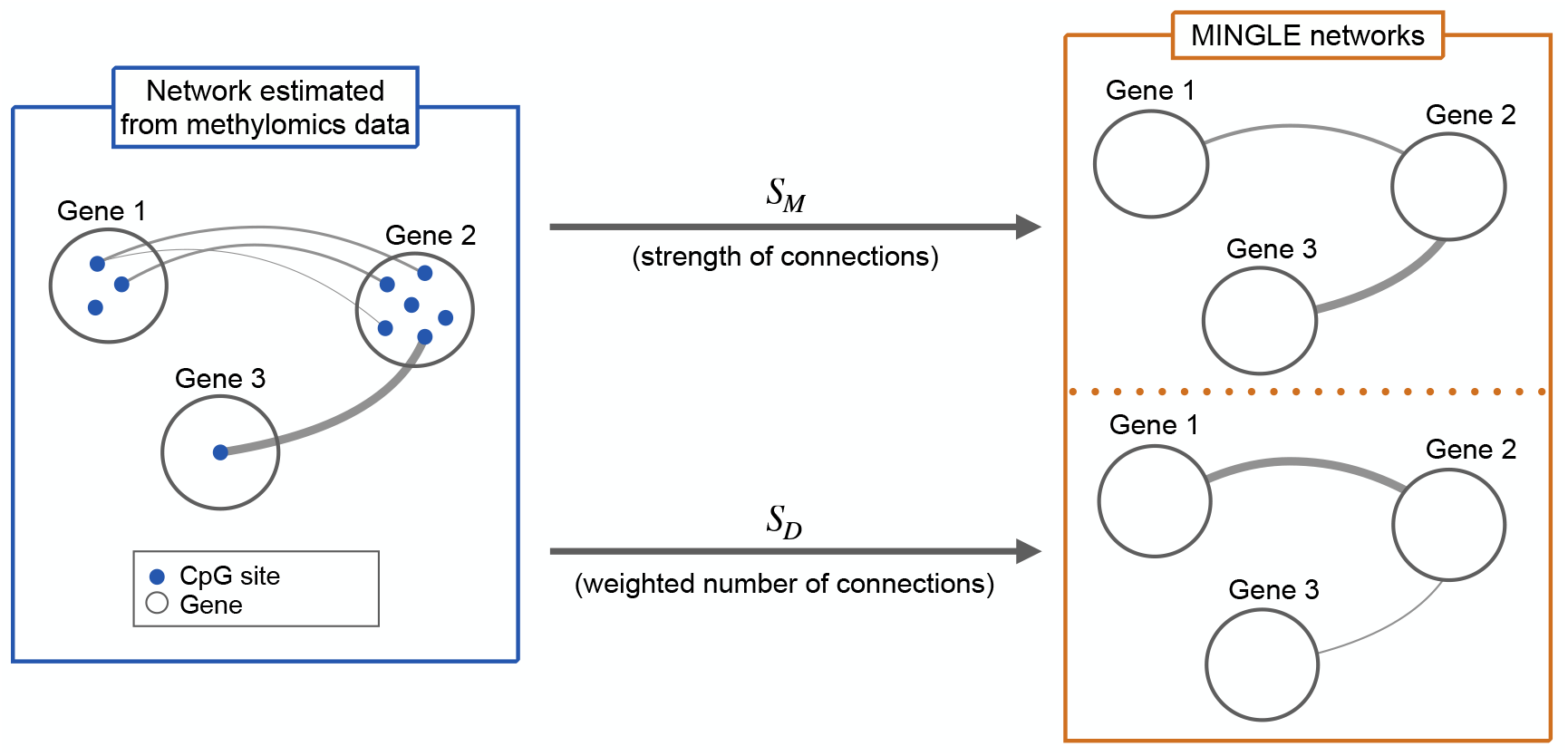
MINGLE conceptual diagram. Scheme representing the transition from the glasso methylomics network (blue box) to MINGLE networks (orange box). The sparse graph estimated from methylomics data is a network of CpG sites (blue nodes). Since each site represents a specific region of a gene (white circles), MINGLE employs this known associations to define networks of genes, whose edges are computed based on the estimated methylomics relations. MINGLE networks can be designed based on the strength of the connections (*SM*, from Eq. (7)), and the number of connections (*DM*, from Eq. (8)). These two definitions lead to networks with different edge weights, while the overall structure of the graphs is the same. Abbreviations: *S*_*M*_, Strength MINGLE network; *D*_*M*_, Degree MINGLE network.

This representation helps the interpretation of results and the identification of important genes at the methylomics level. Once identified the nodes of interest, the original methylomics relations can be recovered to perform a more detailed analysis on the estimated methylomics networks.

## 3. Results and Discussion

In this section, we show and discuss the main results of our integrated multli-omics analysis, with a focus on GBM. We provide the R script to reproduce the analysis, as well as additional tables, which could be independently explored. The list and the corresponding explanation of each additional file is reported in Supplementary Material S1.

### 3.1. Transcriptomics networks

Figure 3a shows the jointly estimated network for the three glioma types, obtained from RNA-Sequencing data with *λ*_1_ = 0.9 and *λ*_2_ = 0.001. Many subnetworks can be clearly distinguished, most of which are small, involving from two to seven nodes. Six big subnetworks are also present (highlighted with green circles in the Figure 3a), which include both exclusive and shared relations among the glioma types. Despite the difficulty of interpretation of such big and dense networks, we observe a prevalence of exclusive edges in some of them. For instance, the green circle A is mainly related to GBM, while B1 and B2 include many oligodendroglioma edges. The small subnetworks are easier to discuss, since they are often related to one glioma type. In particular, when these exclusive relations constitute a subnetwork with more than two nodes, it could represent an important regulation of the associated disease. For example, the grey circle C in Figure 3a is mainly exclusive for GBM, while the two grey circles D1 and D2 include relations of both astrocytoma and oligodendroglioma (which are known to exhibit similarities compared to GBM).

**Figure 3:**
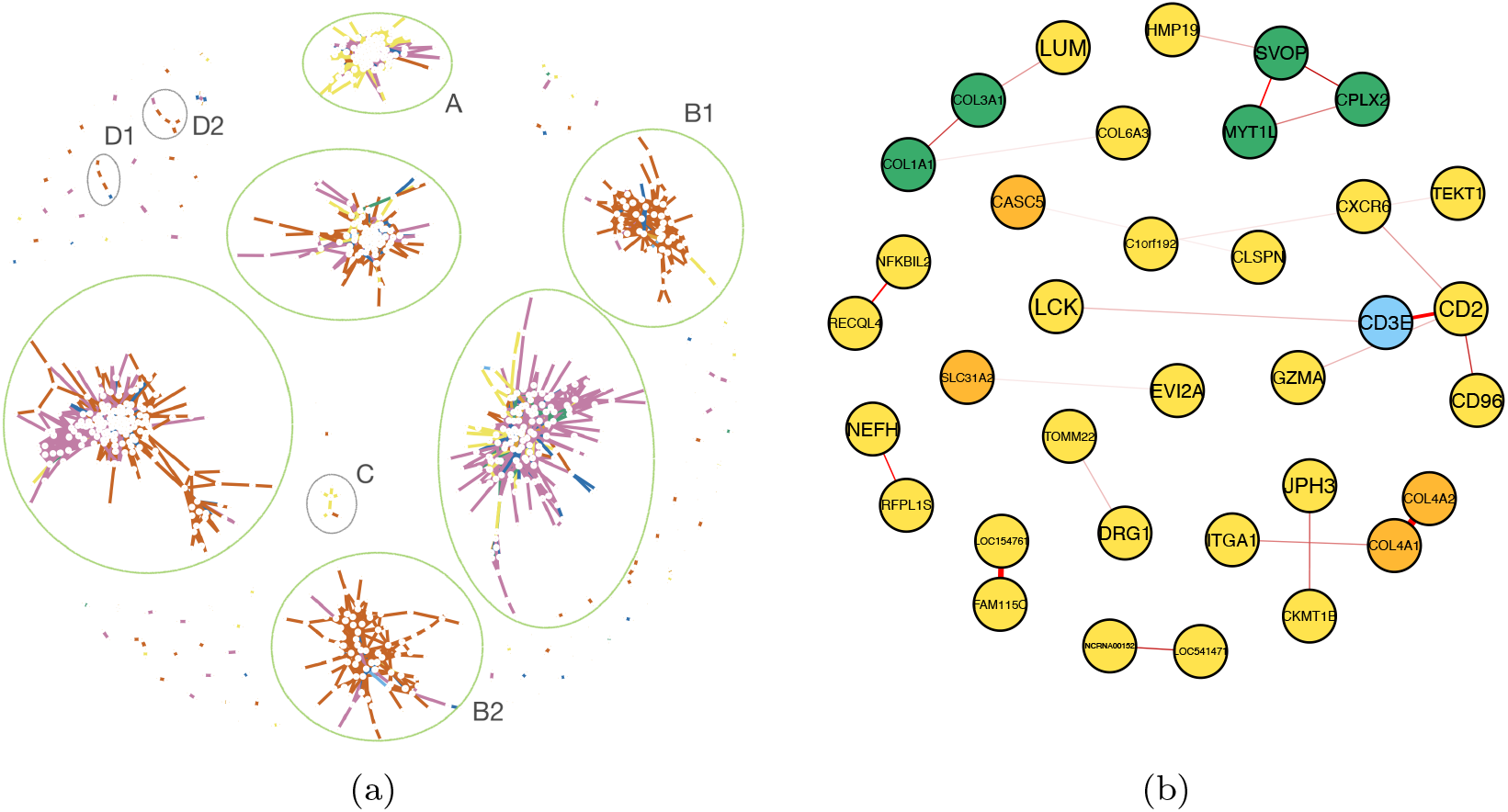
Graphical representations of the estimated networks. (a) Complete JGL networks. This representation is edge-based, i.e., exclusive or shared links are highlighted by different colors. (b) GBM small subnetworks after our filtering process. This representation is node-based, i.e., exclusive or shared genes are highlighted by different colors. 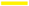 Glioblastoma (GBM) 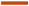 Oligodendroglioma (O) 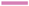 Astrocytoma (A) 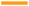 Shared all 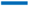 Shared A-O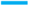 Shared O-GBM 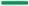 Shared A-GBM.

With the purpose of deeply investigating the GBM glioma type, we focused on the estimated GBM network, which is constituted by 39 subnetworks, divided into 3 big and 36 small subnetworks (less than 6 nodes each). To reduce the complexity of the analysis, we focused on the relations we assume to be more representative of this glioma type, removing the small subnetworks involving only shared genes, or genes of the same family (i.e., trivial links). Our filter removed 23 small subnetworks related either to shared genes or trivial relations, maintaining the relations showed in Figure 3b. By analyzing these subnetworks, we detected some potentially interesting relations, such as:

1. the interaction between *CXCR6, CD2, CD96, CD3E, GZMA* and *LCK* (subnetwork C in Figure 3a). Noteworthy, all these genes are known to be involved in immune-related mechanisms, in some cases explicitly linked to GBM [39, 40, 41, 42, 43, 44].
2. the strong exclusive relations, such as the one between *FAM115C* and *LOC154761*. Both these genes are deferentially regulated in glioma [45, 46], yet *FAM115C* also shows prognostic relevance, as it influences the overall glioma patient’s survival [46].
3. the relations differing with the other glioma types. For instance, in the subnetwork with *COL4A1, COL4A2* and *ITGA1*, the relation between *COL4A1* and *COL4A2* was common to all the glioma types, but only in GBM they are linked with *ITGA1*, which is associated with temozolomide chemotherapy resistance [47]. The biological validation of this estimated link could clarify if the relation among these genes could somehow play a role in GBM treatment resistance.

The interpretation of larger subnetworks might be more challenging, but it could be essential to discover potentially important relations, due to the presence of both mixed and exclusive relations. We report an example of how to accomplish this in the Supplementary Material S3.

### 3.2. MINGLE networks

To further investigate the GBM-exclusive relations by multi-omics integration, we built the corresponding MINGLE networks, i.e., the network of genes based on the GBM-methylomics graph estimated by glasso, defined as described in Section 2.6. For GBM, our results led to the selection of 148 CpG sites, associated with 43 genes. Among them, 23 genes were involved in inter-gene relations, depicted in Figure 4. Interestingly, it might be worthy to note that, differently from the estimated transcriptomics graphs, most of the genes here represented are part of the same network, indicating the presence of a complex structure of gene relations at the methylomics level. Isolated genes are not visualized in these networks, because of their inherent characteristic of having relations solely among their own CpG sites. Despite this, these genes hold relevance as their intra-gene relations might potentially influence GBM processes. The two representations in Figure 4 show the same network with different edge weights, which is represented by edge thickness, allowing the disclosure of distinct but complementary properties of the same relations between CpG sites of pair genes. For instance, *D*_*MG*_ (Figure 4a), which is based on the number of connected probes, shows that the two genes *PLEK* and *PTPN6* are connected by multiple associations at methylomics level. However, these relations are weak, as revealed by the *S*_*MG*_ network, whose edges are computed based on the strength of the connections between the gene-related CpG sites. Conversely, *COL4A3, DOCK2, MAL, FA2H* and *CPLX2* genes appear strongly connected in Figure 4b, while involving few methylomics relations. The proposed MINGLE network representations are very useful to detect genes characterized by potentially relevant methylomics relations, yet loosing the original meaning of the edges as conditional dependence. However, once identified nodes of interest, it is always possible to go back to the original network to further explore the CpG site relations. For instance, Figures 5 and 6 show the estimated networks among the probes of the discussed genes. The two figures provide two different representations, one with the nodes distributed based on their distance in the network (Figures 5a and 6a), which simplify the observation of the overall network structure, and another with CpGs sorted with respect to their location in the genome (Figures 5b and 6b). The latter can be particularly useful to detect if there is a specific region of the gene that appears to be more involved in methylomics relations, which could suggest a specific area particularly accessible to methylation.

**Figure 4:**
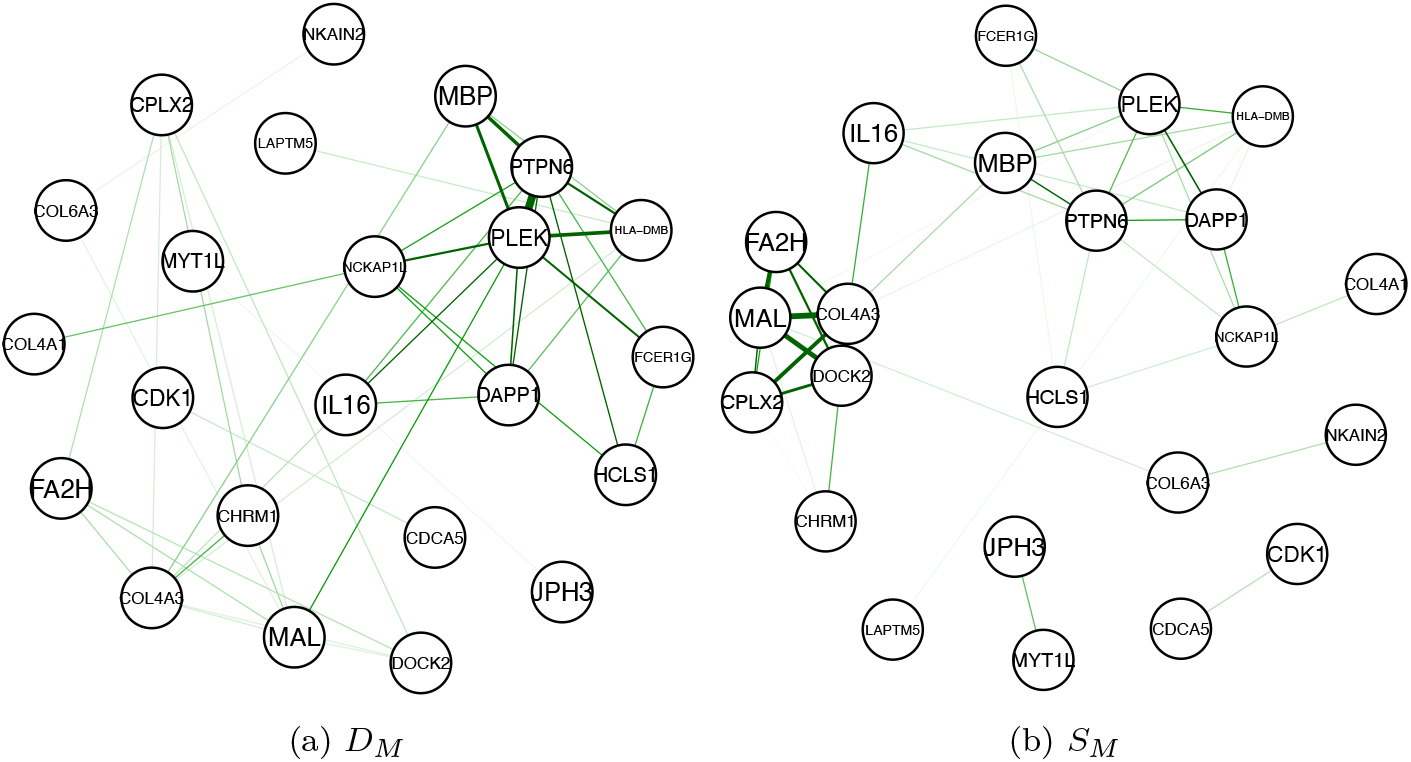
GBM MINGLE networks. The structure of the two graphs is the same (i.e., genes are connected by the same link), while the edge weights are computed by taking into account (a) the number, or (b) the strength of the relations in the methylomics networks. Abbreviations: *S*_*M*_, Strength MINGLE network : *D*_*M*_, Degree MINGLE network.

**Figure 5:**
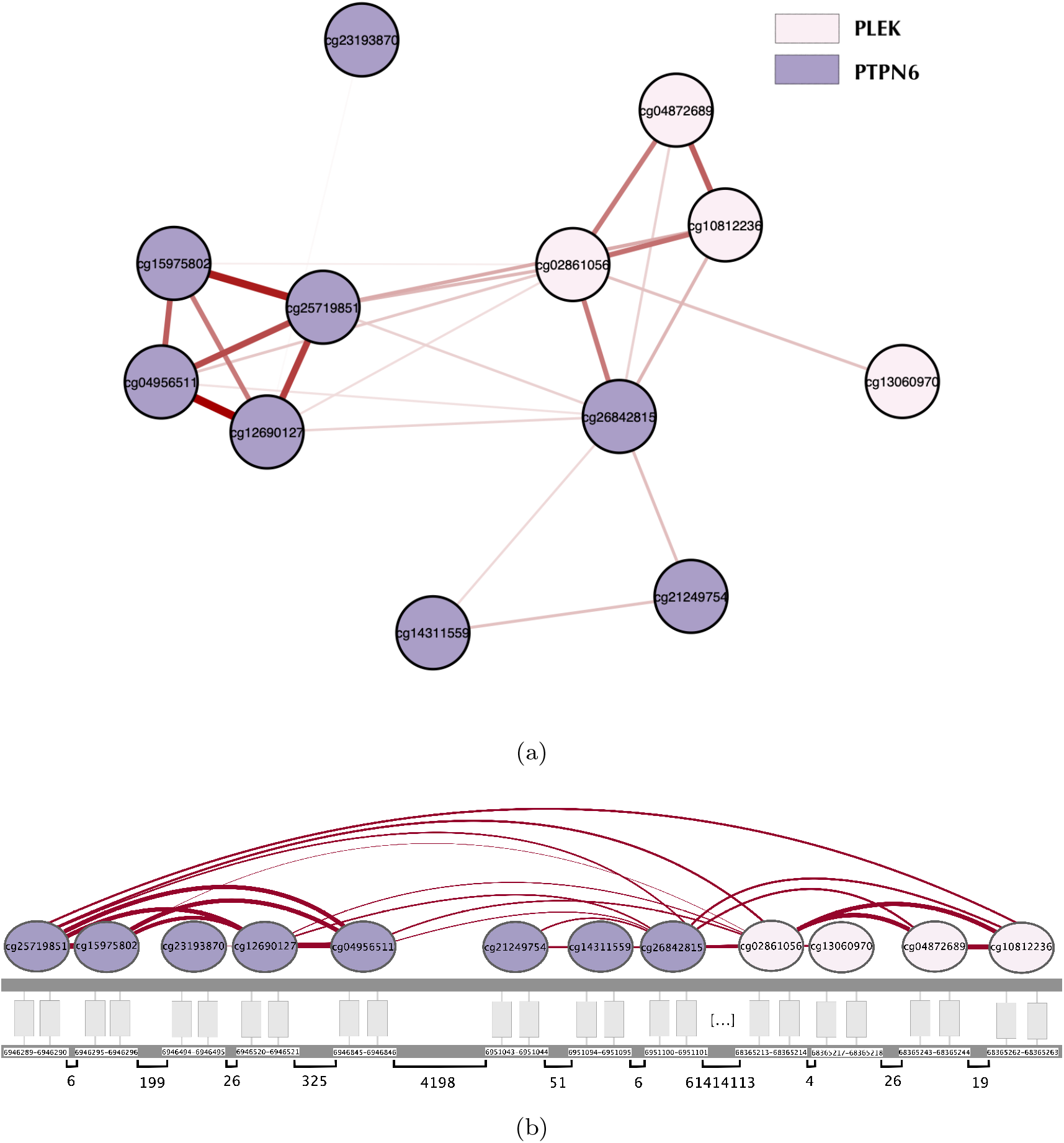
Original estimated relations between CpG sites of the two genes *PLEK* (pink) and *PTPN6* (purple). (a) The network layout is based on the Fruchterman-Reingold force-directed algorithm, which distributes nodes based on their distance in the network (neighbours are closer). (b) The nodes are represented according to their location in the genome. For each CpG, the corresponding nucleotides are reported, along with the computed distance between two subsequent CpGs.

**Figure 6:**
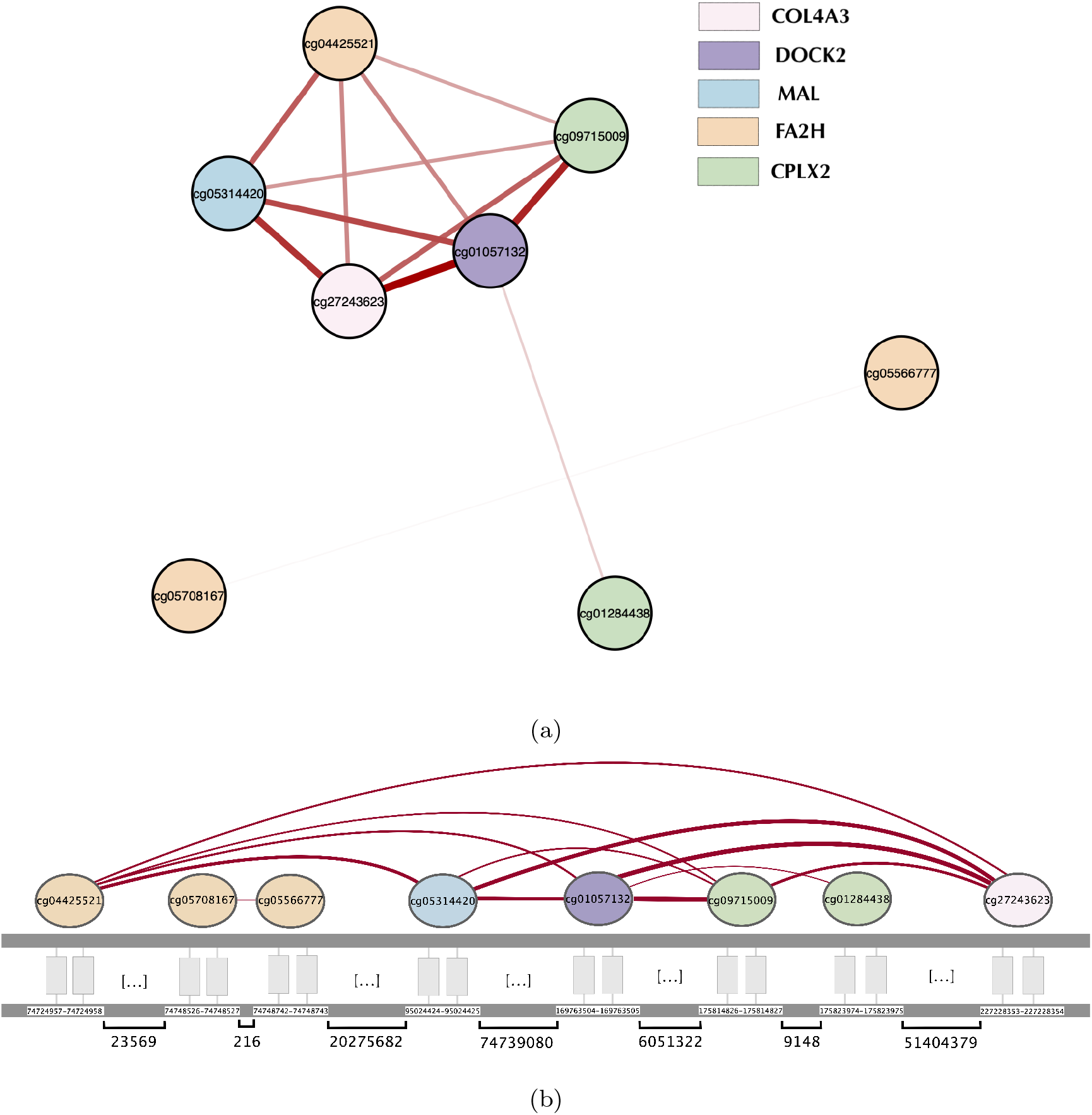
Original estimated relations between CpG sites of the genes *COL4A3* (pink), *DOCK2* (purple), *MAL* (light blule), *FA2H* (orange) and *CPLX2* (green). (a) Network layout is based on the Fruchterman-Reingold force-directed algorithm, which distributes nodes based on their distance in the network (neighbours are closer). (b) The nodes are represented according to their location in the genome. For each CpG, the corresponding nucleotides are reported, along with the computed distance between two subsequent CpGs.

All the information necessary to reproduce similar networks for other variables of interest are collected in the provided Supplementary files.

### 3.3. Classification

As a way to validate the biological significance of the networks inferred, patient classification was performed through sparse multinomial logistic regression.

For transcriptomics, classification algorithm was applied to four datasets, comprising all the glioma samples and: (i) the starting 16655 variables set, (ii) the set of selected variables (union of the variables selected for each glioma type), (iii) variables linked by glioma-type exclusive edges, and (iv) glioma-type exclusive variables (the nodes appearing only in one gliomatype network). By performing classification on these four datasets, we aim to investigate not only if the three glioma types are well characterized by the set of identified genes, but also if exclusive nodes (or exclusive pairs) can provide comparable performances, yet further reducing the number of variables. Moreover, by employing regularized classification, we are able to identify single genes that allow the separation of patients into the different glioma types, thus being promising candidates as diagnostic markers. Regularization can be modulated by the parameter *α*, which was set between 0 (Ridge; no feature selection) and 1 (LASSO; feature selection). Table S5 resumes the classification outcomes for different values of *α*. For *α* = 0, better performances are obtained by considering the complete set of variables. However, despite the considerable variable reduction (*<* 20% of the initial dataset), our identified subsets are also providing excellent performances, with AUC *>* 0.9. Notably, reducing the variables to the exclusive nodes (case (iv)) provides better results, supporting that the genes that are exclusive of one glioma-type network are more biologically representative of the disease. Classification performances enhance with a small regularization.

Regarding methylomics results, we reduced the cases of study to the first two, i.e., starting variables versus selected ones. The other two cases involving exclusive variables were not taken into account, as the methylomics analysis was specifically defined to target CpG sites associated with nodes linked by exclusive relations. The results of classification in the two cases show comparable AUC values for all the values of the regularization parameter (Table S6), proving that the subset of selected CpGs is not loosing the original diagnostic information.

A comprehensive analysis of the classification results for both transcriptomics and methylomics data is reported in the Supplementary Material S4.

### 3.4. Survival Analysis

Survival analysis was performed to detect features having impact on GBM overall survival. To this end, we focused on the final outcome of our multi-omics integration procedure, which selected 148 CpGs associated to 43 genes. We investigated both omics, by including in the survival analysis all the 43 genes, and the top 20 nodes of the methylomics networks, taking into account both, the number of connections, and the median sum of weights (in case of multiple variables with the same ranking, this selection has been expanded to include all the nodes with equivalent importance). Based on transcriptomics data, we detected 27 over 43 genes significantly affecting GBM survival (*p*-value *<* 0.05). Interesting, many of them are isolated in the defined MINGLE networks. A literature research on the genes with lower *p*-value (Figure 7a) revealed that most of them act in cancer-related processes, also in the context of glioma [48, 49, 50, 51]. The only exception regards *PLEK*, about which there are no exhaustive studies investigating its role in cancer. However, based on our estimated networks, this gene is directly linked with *FCER1G* and *PTPN6*, both known to influence glioma patient’ prognosis and treatment responses [52, 53].

**Figure 7:**
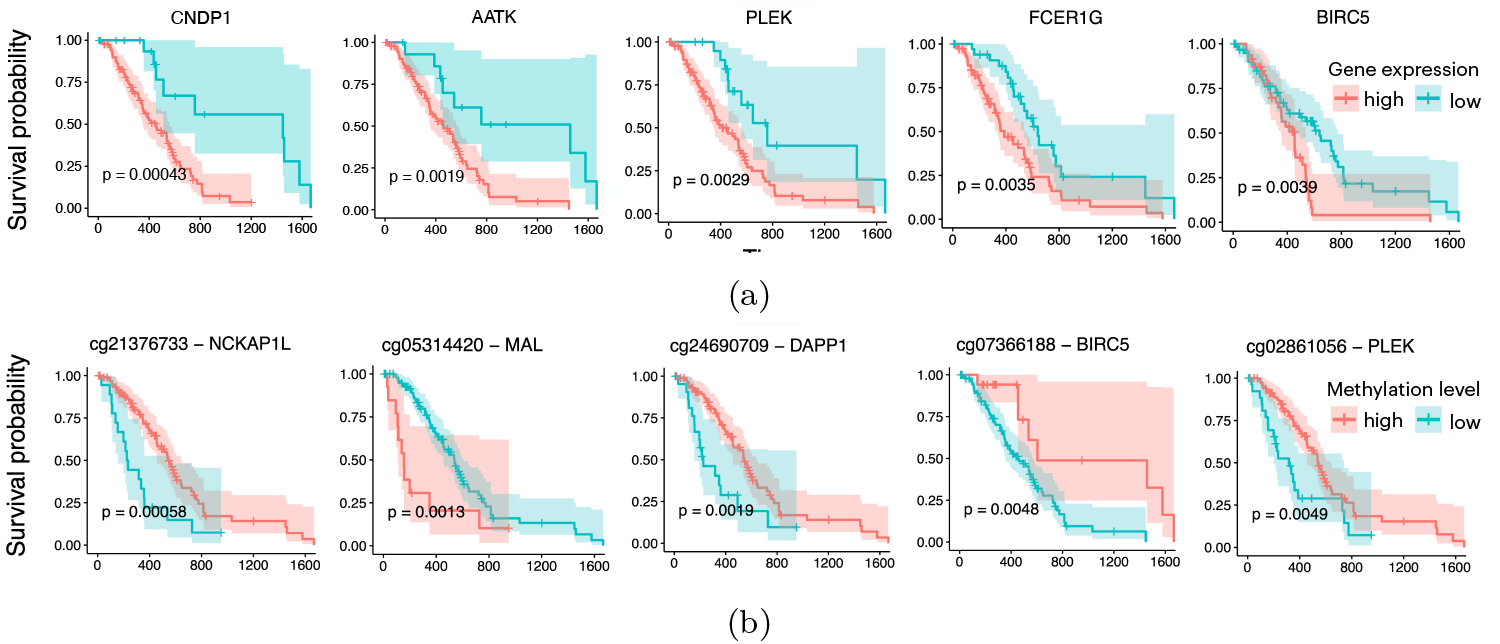
Kaplan-Meier curves of the top 5 (a) genes and (b) CpG sites with lower *p*-values.

In methylomics, a total of 42 variables was investigated, namely 22 highly connected, and 20 strongly connected nodes. Among them, 23 CpGs resulted having impact in GBM prognosis, corresponding to a total of 13 genes. Figure 7b shows the Kaplan-Meier curves for the 5 CpGs with lower *p*-values. Again, we encounter genes already pointed out by previous analyses, such as *MAL, BIRC5*, and *PLEK*. The CpG with the strongest impact on GBM survival is associated to gene *NCKAP1L*, whose probes bind to *PLEK* and *PTPN6* CpG sites (Figure 4), recurrent genes in our study. Although *NCKAP1L* is not yet studied in glioma, it is known to be a tumor suppressor gene [54], and barely expressed in several cancers [54, 55, 56].

Further survival analysis results are available at Supplementary Material S5.

## 4. Conclusion

The presented work represents a pioneering study on multi-omics integration from glioma patients grouped following the latest glioma classification guidelines release. The multi-omics integration through MINGLE led to the identification of gene subnetworks that could be associated with the specific glioma-type. The two omics data revealed different network structures, as we can see some connections among CpGs of genes that were not related in RNA Sequencing-based networks. This result further stresses the importance of a multi-omics study, which allows the disclosure of unknown mechanisms that would be impossible to detect by a single-omics analyis.

Patient classification provided comparable results when considering the complete dataset and our subset of identified features, in both omics. In particular, the exclusively selected variables were recognized, among the whole subset of detected genes, as the ones that better distinguish the patients belonging to different glioma types. The diagnostic variables selected via model regularization from the original starting dataset (case (i)) are different from the ones selected from the reduced set (cases (ii)-(iv)), indicating the existence of different sets of genes suitable for classification purposes. Nevertheless, the proposed approach also estimates a network structure, which represents a remarkable advantage in terms of biological interpretability, being extremely important for a better understanding of the glioma molecular mechanisms.

Survival analysis focused on GBM patient stratification revealed that around 50% of the investigated variables have prognostic value, pointing out potential interesting biomarkers from a multi-omics perspective, such as *AATK, PLEK, PTPN6*, and *BIRC5*. In particular, most of these genes, which are involved in GBM-exclusive relation from the transcriptomics layer, appear as isolated nodes in the MINGLE network, suggesting that they are solely characterized by links among their own CpG sites. If these findings are biologically verified, these genes could hold significant promise as therapeutic targets for GBM. Indeed, despite their relevance to the disease, the predicted limited impact of such genes on the overall network suggests that targeted modifications may increase the specificity and decrease effects in other other crucial regulatory processes.

The proposed framework could be generalized to the cases in which merging network information coming from different omics could be useful for biological interpretation purposes. For instance, MINGLE networks could be designed from other omics with previously known associations, such as transcriptomics and proteomics, or genomics. Also, MINGLE could be applied to other data types and contexts, as a general tool for integrating multi-layers networks when the functional link between the variables of two layers is available.

Furthermore, this work provides customizable tables that serve as a tool to independently explore the obtained results. These tables collect genes and CpG-site relations, specific for each glioma type, resulting from our network inference, classification, and survival outcomes. The exploration of such files might lead to the detection of other promising groups of features as novel potential biomarkers or worth of future laboratory validation. These results might motivate further investigation on the role of the genes we identified as potential biomarkers, with the aim of improving the clinical management of glioma patients.

## CRediT author statement

**Roberta Coletti:** Conceptualization, Methodology, Software, Validation, Formal analysis, Writing - Original Draft, Visualization.

**João Carrilho:** Methodology, Formal analysis, Software (classification).

**Eduarda P. Martins:** Conceptualisation, Writing - Review and Editing.

**Céline S. Goçalves:** Conceptualisation, Writing - Review and Editing.

**Bruno M. Costa:** Conceptualisation, Writing - Review and Editing, Funding acquisition.

**Marta B. Lopes:** Conceptualization, Methodology, Supervision, Writing - Review and Editing, Funding acquisition.

## Acknowledgements

This work was supported by national funds through Fundaç ão para a Ciê ncia e a Tecnologia (FCT) with references CEECINST/00042/2021, 2021.02600.CEECIND (doi: 10.54499/2021.02600.CEECIND/CP1664/CT0015), UMINHO/BIM-CNCG/2023/102, UIDB/00297/2020 and UIDP/00297/2020 (NOVA Math, Center for Mathematics and Applications), UIDB/00667/2020 and UIDP/00667/2020 (UNIDEMI), CEECIND/00072/2018/CP1581/CT0011 (doi: 10.54499/CEECIND/00072/2018/CP1581/CT0011), CEECINST/00077/2018/CP1640/CT0002 (doi: 10.54499/CEECINST/00077/2018/CP1640/CT0002), 2022.04859.PTDC (doi: 10.54499/2022.04859.PTDC), and the research project “MONET – Multi-omic networks in gliomas” (PTDC/CCI-BIO/4180/2020). This work was also supported by the project NORTE-01-0145-FEDER-000055, supported by Norte Portugal Regional Operational Programme (NORTE 2020), under the PORTUGAL 2020 Partnership Agreement, through the European Regional Development Fund (ERDF).

