## Supplementary Materials for "A Novel Tool for Multi-Omics Network Integration and Visualization: A Study of Glioma Heterogeneity"

### S1. Supplementary files and contents

This work includes additional tables which can be independently explored to identify variables or relations of interest, as potential glioma biomarkers. We provide a total of four tables, described in the following.

- *Transcriptomic-Joint-Network*: matrix representing the joint network estimated from transcriptomics data. Rows and columns list the union of the genes involved in relations in the three glioma types. Cells report either 0, meaning that no links exist between the corresponding genes, or a label specifying the glioma type for which the relation is estimated. If two genes are connected, the cell is colored depending on the cancer type involved, according to the following rules: yellow (GBM), red (oligodendroglioma), pink (astrocytoma), blue (astrocytoma - oligodendroglioma), light blue (oligodendroglioma - GBM), green (astrocytoma - GBM), and orange (shared by all).

- *Methylomic-Joint-Network*: matrix representing the joint network estimated from transcriptomics data. Even if in this case the networks are estimated separately, we joined the information to facilitate the exploration of the results. In this file there are two sheets. Sheet1: rows and columns list the union of the CpG sites involved in relations in the three glioma types. Sheet2: rows and columns list the genes associated to the CpG sites selected by methylomics layer. It corresponds to the joint representation of the MINGLE networks (Section 2.5). We note that in this matrix there could be isolated nodes, i.e., genes with no links within the MINGLE network, meaning that it is selected due to inner-gene methylomics relations. In both sheets, cells report 0 or a label and a color associated to the corresponding glioma type for which the relation is estimated, according to the following rules: yellow (GBM), red (oligodendroglioma), pink (astrocytoma), blue (astrocytoma - oligodendroglioma), light blue (oligodendroglioma - GBM), green (astrocytoma - GBM), and orange (shared by all).

- *Table1-Integrated_multi-omics_results*: table joining the results obtained by our multi-omics analysis. The file contains two sheets. Sheet1: the selected CpG sites are associated to the genes, specifying the location of each probe on the genome. "Methylomics - glioma type" columns report the glioma type for which the CpG has been selected. The table also highlights in green the genes involved only in intragene methylomics relations in the MINGLE networks, by specifying the corresponging glioma type. Additionally, the "Transcriptomics - glioma type" column reports the glioma types in which the genes are connected, allowing the comparison between the two-omics results. Sheet2: for each gene, the table reports the total CpG sites in our dataset, and the percentage of the selected probes after glasso algorithm. These values are used to compute the edges of $D_{MG}$ network according to Equation (8).

- *Table2-Classification&Survival*: resumes the classification and survival results in three sheets. Sheet1: focused on transcriptomics data, it reports the list of the genes detected by classification ($\alpha = 0.1$) on the subset of variables selected by our network-based approach. Each gene is associated to one or more glioma type by the classification algorithm, as specified by the columns designed as "Classification - glioma type". Columns named "Network - glioma type" report if the gene is present in a specific glioma-type network, and if it is involved in exclusive relations. The last column reports the result (in term of $p$-value) of the subset of genes investigated by survival analysis (see Section S5 for details). The acronym *NE* stands for "Not Evaluated", and labels the genes that were not included in this analysis. Sheet2: focused on methylomics data, it reports the list of the CpG sites detected by classification ($\alpha = 0.1$) on the subset of variables selected by our network-based approach. Each probe is linked with the corresponding gene, and the glioma types for which it was selected by both network inference and classification methods. The table also highlights in blue the genes detected by classification on transcriptomics, and, in positive case, the associated glioma type is included. The last column reports the result (in term of $p$-value) of the subset of CpG sites investigated by survival analysis (see Section S5 for details). The acronym *NE* stands for "Not Evaluated", and labels the probes that were not included in this analysis. Sheet3: reporting survival analysis results presented in the main text, i.e., focused on CpG sites and associated genes derived from our two-steps network-based variable selection framework. The investigated CpG sites and corresponding genes are listed in the first two columns. The subsequent two columns report the $p$-values of the statistically significant variables, for both methylomics (CpG.pvalue) and transcriptomics (Gene.pvalue). Note that genes may be duplicated if more than one of their CpG sites is analyzed. In these cases, the $p$-value is reported at the gene's first occurrence only.

These contents will be available after publication.

### S2. Network regularization parameter selection and mathematical validation

To estimate networks by glasso or JGL, a set of parameters is needed to regulate network sparsity. Although several tools to drive the parameter selection process are available, the estimated optimal values are usually not suitable for such high-dimensional problems, as they lead to too dense or not informative networks. For instance, by taking into account the Extended Bayesian Information Criterion (EBIC) [1], which is particularly suitable in case of high dimensional models, the estimated optimal JGL networks would be constituted by only four nodes, and two edges, shared across the three glioma types. For this reason, we followed the suggestion of the authors of JGL, who recommend to set regularization based on practical consideration, such as the overall network interpretability [2].

In our work, JGL is applied to transcriptomics data. The penalty function is regulated by two parameters, controlling the amount of sparsity ($\lambda_1$), and the similarity across the classes ($\lambda_2$). In this case, our goal was to explore the differences among the three glioma types without forcing edges to be the same, thus we tested low values of $\lambda_2 \in \{0.001, 0.01, 0.1\}$. Conversely, we set high $\lambda_1$ values ($\lambda_1 \in \{0.85, 0.90, 0.95, 0.97\}$), to reduce the set of selected variables to an interpretable amount of node in each graph. Based on the results obtained for different parameter settings (Table S1), we excluded the extreme values due to either computational problems or not suitable outcomes. In particular, by setting $\lambda_1 = 0.85$, the algorithm does not converge due to the high dimension of the dataset, while $\lambda_1 = 0.97$ determines too strong variable reduction. On the other hand, $\lambda_2 = 0.1$ leads to shared features only, thus not informative for our purposes. We focused our analysis on the results obtained by setting $\lambda_1 = 0.9$, which led to a network sufficiently complex and sparse to be informative and interpretable, while reducing the similarity across classes ($\lambda_2 = 0.001$).

The second step of the proposed pipeline was based on mehtylomics data. As our integrative multi-omics analysis only focuses on the CpG sites associated to genes that were linked by exclusive edges in transcriptomics networks, we consider each glioma type per time. This simplification allows the employment of the classical glasso, which penalty is regulated by a single parameter $\rho$. We tested several regularization parameters, yet we fixed $\rho = 0.85$, as it induces a strong variable reduction while maintaining a complex network structure (Table S2). Also, in the particular case of GBM (the glioma type on which this study is focused), this parameter setting also lead to the optimal glasso model according to the EBIC criteria [1].

Table S1: Resume of JGL results for different combination of the tuning parameters $\lambda_1 \in \{0.90, 0.95, 0.97\}$ and $\lambda_2 \in \{0.001, 0.01, 0.1\}$. The case of $\lambda_1 = 0.85$ is not shown since the algorithm does not converge. For each combination, the number of nodes (genes) and edges per glioma type are reported. The values in brackets indicate the exclusive counts, i.e., the number of nodes or edges unique to each glioma-type-specific network. The last column shows the number of edges shared across the three glioma types. Astro: astrocytoma; Oligo: oligodendroglioma; GBM: glioblastoma.

| $\lambda_1$ | $\lambda_2$ | **Nodes** | | | **Edges** | | | **Shared edges** |
|---|---|---|---|---|---|---|---|---|
| | | Astro | Oligo | GBM | Astro | Oligo | GBM | |
| 0.90 | 0.1 | 258 (0) | 258 (0) | 258 (0) | 913 (0) | 913 (0) | 913 (0) | 913 |
| 0.90 | 0.01 | 384 (41) | 422 (109) | 261 (31) | 1870 (411) | 1790 (405) | 719 (57) | 576 |
| 0.90 | 0.001 | 517 (116) | 611 (254) | 300 (71) | 3026 (1502) | 2940 (1540) | 782 (303) | 297 |
| 0.95 | 0.1 | 25 (0) | 25 (0) | 25 (0) | 27 (0) | 27 (0) | 27 (0) | 27 |
| 0.95 | 0.01 | 39 (0) | 37 (0) | 25 (0) | 38 (0) | 36 (1) | 24 (0) | 23 |
| 0.95 | 0.001 | 87 (33) | 75 (22) | 24 (5) | 131 (74) | 102 (48) | 19 (4) | 8 |
| 0.97 | 0.1 | 4 (0) | 4 (0) | 4 (0) | 2 (0) | 2 (0) | 2 (0) | 2 |
| 0.97 | 0.01 | 4 (0) | 4 (0) | 4 (0) | 2 (0) | 2 (0) | 2 (0) | 2 |
| 0.97 | 0.001 | 7 (0) | 8 (1) | 4 (0) | 5 (0) | 6 (1) | 2 (0) | 2 |

Table S2: Resume of glasso results for different combination of the tuning parameter $\rho \in \{0.75, 0.80, 0.90, 0.95\}$ . For each parameter value, the number of nodes (CpGs) and edges per glioma type are reported. The last column shows the number of the genes corresponding to the selected CpGs. Astro: astrocytoma; Oligo: oligodendroglioma; GBM: glioblastoma.

| $\rho$ | **Nodes** | | | **Edges** | | | **Corresponding genes** | | |
|---|---|---|---|---|---|---|---|---|---|
| | Astro | Oligo | GBM | Astro | Oligo | GBM | Astro | Oligo | GBM |
| 0.75 | 1416 | 1672 | 499 | 10572 | 26267 | 1180 | 302 | 366 | 123 |
| 0.80 | 783 | 1178 | 283 | 4391 | 14458 | 532 | 213 | 303 | 75 |
| 0.85 | 413 | 685 | 148 | 1242 | 5658 | 179 | 124 | 219 | 43 |
| 0.90 | 142 | 278 | 66 | 145 | 1162 | 58 | 48 | 102 | 22 |
| 0.95 | 15 | 52 | 14 | 9 | 49 | 7 | 7 | 21 | 7 |

To validate the reliability of the estimated networks, we would show that the set of variables selected by the algorithm is the one leading to the best solution of the optimization problem in equation (1), i.e., no other combinations of the same number of variables leads to comparable performances. To this aim, we compared the objective functions related to the optimal set of variables $D^*$, with the ones computed by considering 1000 random sets of variables such that $\dim(D_i) = \dim(D^*)$, for all $i = 1, ...1000$. We considered the objective function of glasso (Eq. (1)) as a difference between two terms: the main function $O(\Theta) = \log(\det(\Theta)) - tr(S\Theta)$, and the penalty $P(\Theta) = \rho||\Theta||_1$. By fixing $\rho \sim 0$, the optimization problem is driven by the first term, as $O(\Theta) >> P(\Theta)$. Let $\Theta^*$ be the precision matrix for the optimal set $D^*$, obtained by imposing a low value of $\rho$ in order to avoid further variable reduction. Let $\Theta^{D_i}$ be the precision matrix estimated for the random dataset $D_i$, with the same regularization parameter. If $O(\Theta^*) >> O(\Theta^{D_i})$, for all $i = 1, ..., 1000$, we can conclude that there are no multiple sets of variables leading to the optimal solution.

Note that the same validation approach was applied for validating both transcriptomics and methylomics results, despite transcriptomics networks being estimated by the JGL method, which defines a slightly different objective function. However, adapting the mathematical validation to the JGL algorithm is not suitable as the outcomes could be affected by a compensation bias, which would alter the optimal solution, making it impossible the comparison with the random sets of variables. Given that the objective functions of the two methods mainly differ for the penalty term, the decision of considering glasso for both omics network validation is reasonable if the order of magnitude of the penalty is much smaller than the one of the main term $O(\Theta)$. In this way, the JGL objective function can be considered as a sum of $O(\Theta^{(l)})$, for $l = 1, 2, 3$, and the robustness of the optimization algorithm can be proved for each glioma type, separately, avoiding the potential compensation bias. The results of the performed validation are reported in Tables S4 and S3.

Table S3: Mathematical validation of transcriptomics results, for the different glioma types. $O(\Theta^*)$ and $P(\Theta^*)$ are, respectively, the main function and the penalty computed by considering the optimal set of variables. $O(\Theta^D)$ and $P(\Theta^D)$ summarize the results obtained for random variable subsets by showing the best, the average and median values computed. Table also shows in the first column the values of the regularization parameter $\rho$ fixed in each case. Astro: astrocytoma; Oligo: oligodendroglioma; GBM: glioblastoma.

| | $\rho$ | $O(\Theta^*)$ | $O(\Theta^D)$ | | | $P(\Theta^*)$ | $P(\Theta^D)$ | | |
|---|---|---|---|---|---|---|---|---|---|
| | | | Best | Average | Median | | Best | Average | Median |
| Astro | 0.07 | 481.03 | 37.68 | 14.40 | 14.26 | 0.85 | 0.75 | 0.71 | 0.71 |
| Oligo | 0.1 | 486.824 | 53.24 | 28.91 | 29.12 | 0.88 | 0.85 | 0.78 | 0.78 |
| GBM | 0.009 | 536.564 | 350.31 | 339.97 | 340.10 | 0.61 | 0.61 | 0.61 | 0.61 |

Table S4: Mathematical validation of methylomics results for the different glioma types. $O(\Theta^*)$ and $P(\Theta^*)$ are, respectively, the main term and the penalty computed by considering the optimal set of variables. $O(\Theta^D)$ and $P(\Theta^D)$ summarize the results obtained for random variable subsets by showing the best, the average and median values computed. Table also shows in the first column the values of the regularization parameter $\rho$ fixed in each case. Astro: astrocytoma; Oligo: oligodendroglioma; GBM: glioblastoma.

| | $\rho$ | $O(\Theta^*)$ | $O(\Theta^D)$ | | | $P(\Theta^*)$ | $P(\Theta^D)$ | | |
|---|---|---|---|---|---|---|---|---|---|
| | | | Best | Average | Median | | Best | Average | Median |
| Astro | 0.01 | 436.08 | 184.28 | 166.65 | 166.61 | 0.46 | 0.50 | 0.45 | 0.45 |
| Oligo | 0.08 | 366.53 | -76.82 | -76.96 | -56.36 | 0.81 | 0.74 | 0.74 | 0.74 |
| GBM | 0.001 | 227.96 | 130.05 | 120.38 | 120.39 | 0.38 | 0.44 | 0.36 | 0.36 |

### S3. Further analysis of transcriptomics networks

The GBM estimated transcriptomics network is composed by many subnetworks, as introduced in Section S5. Among the three big subnetworks, we considered only the one highlighted in Figure 3a by the green circle A, which is mainly constituted by exclusive GBM relations. Most of the nodes of this network (Figure S1) are involved in relations that are common to all the glioma types. However, among them, some genes exclusively selected for GBM appear, suggesting the correspondent relations as relevant for GBM only. As an example, *SGOL2* and *ECT2* (yellow nodes in the right side of the figure) are connected with *BUB1*, which is a key gene in the GBM network given the high number of connections. *BUB1* is known to be involved in several glioma processes, such as proliferation, migration, and infiltration, and it is overexpressed in glioma patients with poor prognosis [3]. In GBM, the protein coded by *ECT2* is associated with tumor invasiveness [4]. Even if its role in glioma has not yet linked to *BUB1*, in gastric cancer it has been shown that *ECT2* upregulation is correlated with the co-overexpression of both *E2F7* and *BUB1*, genes contributing to the transcription of cancer stem cells [5], making it a promising candidate for further studies.

### S4. Classification results

Regularized classification was performed by taking into account datasets composed by the same samples but different sets of variables, from both omics. In particular, for transcriptomics, four variable sets were considered: (i) the starting 16655 variables (ii) the set of selected variables (union of the ones selected for each glioma type), (iii) variables linked by glioma-type exclusive edges, and (iv) glioma-type exclusive variables (the nodes appearing only in one glioma-type network). For methylomics, we reduced

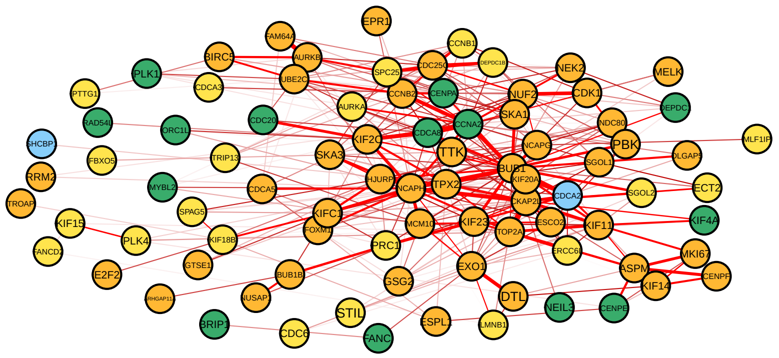


Figure S1: Big GBM subnetwork A in Figure 3a in a node-central representation. The colour layout used for this network representation is coherent with the one presented with the joint networks in Figure 3a. Specifically, Glioblastoma (GBM) Oligodendroglioma (O) Astrocytoma (A) Shared all Shared A-O Shared O-GBM Shared A-GBM .

our analysis to the first two cases, i.e., the starting dataset versus a reduced subset of selected variables. The complete results obtained by varying $\alpha$ in [0, 1] are reported in Tables S5 and S6.

Considering both omics, the identified sets of variables provide good results in term of AUC values, which were comparable for different $\alpha$ values. In transcriptomics (Table S5), for $\alpha = 0$, the set of exclusive variables (Ex.Nod.) leads to better AUC values with less number of selected variables, further supporting the importance of the exclusive genes in distinguish the three glioma types. The Venn diagram (figure S2) also confirms the relevance of exclusiveness for classification purposes, since most of the features identified by the regularized classification method are either exclusive variables or related to exclusive edges, independently from the regularization strength. By analyzing the subset of features selected by the classifier on transcriptomics, we discovered three variables commonly detected in all four cases (ALL, ALL-Sel., Ex.Edg. and Ex.Nod.), i.e., the genes *RAB36*, *RPL22*, and *CHGB*. Independently by the value of $\alpha$, most of the genes detected by starting from the complete dataset (case ALL) are different from the ones coming from our framework, highlighting that there could be more than one set of features impacting on glioma-type classification.

By comparing transcriptomics and methylomics variable selection, there are some probes selected by the classifier that are linked to genes also detected by transcriptomics layer. By fixing $\alpha = 0.1$, our results identi-

Table S5: Classification results from transcriptomics depending on the parameter value $\alpha$. For each case of study, it is shown the number of selected variables, and the AUC values for Train and Test sets. ALL: complete dataset; Sel.: dataset constituted by the variables selected by JGL; Ex.Edg.: dataset constituted by the variables linked by edges exclusive of one glioma-type.; Ex.Nod.: dataset constituted by the variables exclusively selected for one glioma-type.

| $\alpha$ | **n. variables** | | | | **AUC − Train** | | | | **AUC − Test** | | | |
|---|---|---|---|---|---|---|---|---|---|---|---|---|
| | ALL | Sel. | Ex.Edg. | Ex.Nod. | ALL | Sel. | Ex.Edg. | Ex.Nod. | ALL | Sel. | Ex.Edg. | Ex.Nod. |
| 0.0 | 16992 | 853 | 810 | 441 | 0.994 | 0.961 | 0.957 | 0.962 | 0.989 | 0.945 | 0.942 | 0.955 |
| 0.1 | 294 | 169 | 167 | 136 | 0.992 | 0.992 | 0.992 | 0.991 | 0.988 | 0.983 | 0.982 | 0.983 |
| 0.2 | 147 | 107 | 102 | 89 | 0.993 | 0.991 | 0.992 | 0.991 | 0.989 | 0.984 | 0.982 | 0.983 |
| 0.3 | 92 | 81 | 78 | 70 | 0.993 | 0.992 | 0.992 | 0.991 | 0.988 | 0.984 | 0.983 | 0.983 |
| 0.4 | 60 | 68 | 66 | 56 | 0.993 | 0.991 | 0.992 | 0.991 | 0.989 | 0.983 | 0.983 | 0.983 |
| 0.5 | 47 | 60 | 54 | 46 | 0.993 | 0.992 | 0.992 | 0.991 | 0.989 | 0.983 | 0.982 | 0.982 |
| 0.6 | 34 | 48 | 45 | 39 | 0.993 | 0.993 | 0.992 | 0.991 | 0.989 | 0.982 | 0.981 | 0.982 |
| 0.7 | 25 | 37 | 35 | 30 | 0.993 | 0.993 | 0.992 | 0.992 | 0.990 | 0.982 | 0.981 | 0.982 |
| 0.8 | 21 | 30 | 27 | 28 | 0.993 | 0.993 | 0.992 | 0.992 | 0.990 | 0.982 | 0.981 | 0.981 |
| 0.9 | 19 | 23 | 23 | 25 | 0.993 | 0.993 | 0.991 | 0.991 | 0.989 | 0.980 | 0.979 | 0.980 |
| 1.0 | 14 | 18 | 17 | 20 | 0.993 | 0.993 | 0.990 | 0.990 | 0.988 | 0.979 | 0.978 | 0.978 |

Table S6: Classification results from methylomics depending on the parameter value $\alpha$. For each case of study, it is shown the number of selected variables, and the AUC values for Train and Test sets. Start.DS: starting dataset; Sel.: dataset constituted by the subset of variables selected by glasso.

| $\alpha$ | **n. variables** | | **AUC − Train** | | **AUC − Test** | |
|---|---|---|---|---|---|---|
| | Start.DS | Sel. | Start.DS | Sel. | Start.DS | Sel. |
| 0.0 | 11724 | 1045 | 0.994 | 0.984 | 0.985 | 0.977 |
| 0.1 | 248 | 224 | 0.994 | 0.992 | 0.991 | 0.989 |
| 0.2 | 126 | 141 | 0.994 | 0.993 | 0.991 | 0.989 |
| 0.3 | 91 | 92 | 0.993 | 0.993 | 0.991 | 0.989 |
| 0.4 | 64 | 76 | 0.993 | 0.993 | 0.991 | 0.989 |
| 0.5 | 54 | 61 | 0.993 | 0.993 | 0.991 | 0.988 |
| 0.6 | 41 | 49 | 0.993 | 0.993 | 0.990 | 0.988 |
| 0.7 | 35 | 41 | 0.993 | 0.993 | 0.990 | 0.988 |
| 0.8 | 27 | 27 | 0.993 | 0.993 | 0.988 | 0.988 |
| 0.9 | 17 | 24 | 0.993 | 0.993 | 0.986 | 0.987 |
| 1.0 | 11 | 21 | 0.993 | 0.993 | 0.986 | 0.986 |

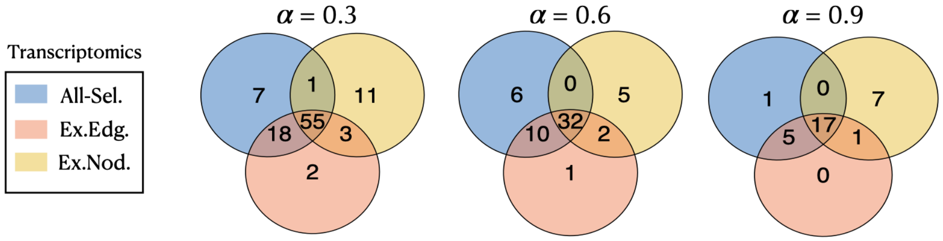


Figure S2: Venn diagrams showing the number of genes selected by regularized classification with different values of $\alpha$. The three sets corresponds to the cases of study involving the genes selected by the network-based approach.

fied 28 common genes. To explore these results, we refer to the *Table2-Classification&Survival* file, Sheets 1 and 2.

### S5. Additional survival analysis

In addition to the survival analysis performed as a validation of glasso outcomes, we also evaluated the prognostic value of the variables that consistently appeared relevant for GBM across various perspectives of our analysis. In particular, we selected a subset of features that were associated to GBM by the classifier, and being present in the corresponding GBM network estimated based on transcriptomics and methylomics. In both omics, 46% of the evaluated genes or CpG sites resulted as statistically significant for GBM survival (p-value $< 0.05$), allowing the distinction of the patients into high- and low-risk groups. In the case of methylomics, by associating the genes to the detected probes, we observed that, over 12 CpG sites with prognostic value, four are linked to *ALOX5* gene, while three are CpGs of *PARVG* gene. Figure S3 shows the Kaplan-Meier curves of the four genes and CpGs with lowest p-value, and therefore being very promising as potential biomarkers. We note that three over four of the most promising CpG sites are probes of *ALOX5* gene, which is known to increase cancer aggressiveness in several cancers [6, 7, 8]. Recently, the role of *ALOX5* in glioma has been studied, revealing its overexpression in cancer compared to normal tissues [9, 10]. Moreover, *ALOX5* potentially affects the regulation of tumor immune-cells infiltration in LGG, and it negatively correlates with treatment sensitivity in both LGG and GBM.

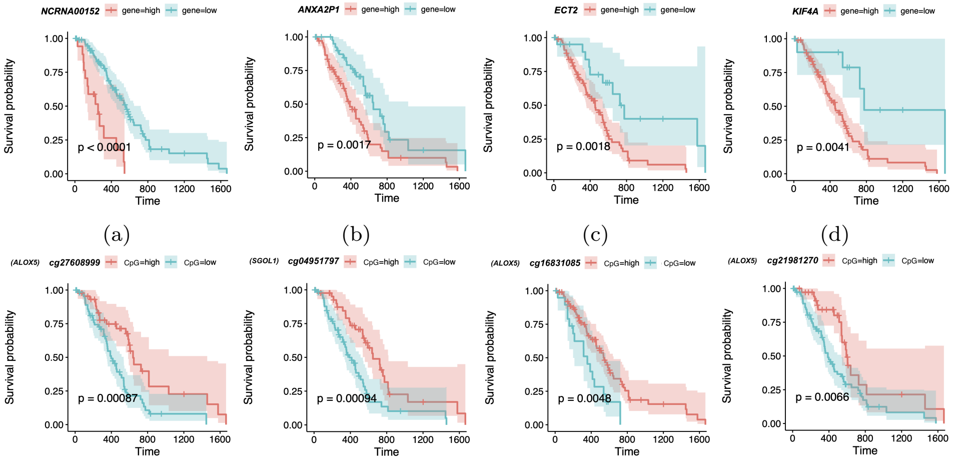


(a) (b) (c) (d)

(e) (f) (g) (h)

Figure S3: Kaplan-Meier curves.(a-d) Transcriptomics; (e-h) Methylomics.
